# Early-life social environment alters juvenile behavior and neuroendocrine function in a highly social cichlid fish

**DOI:** 10.1101/281097

**Authors:** Tessa K. Solomon-Lane, Hans A. Hofmann

## Abstract

Early-life experiences can shape adult behavior, with consequences for fitness and health, yet fundamental questions remain unanswered about how early-life social experiences are translated into variation in brain and behavior. The African cichlid fish *Astatotilapia burtoni*, a model system in social neuroscience, is well known for its highly plastic social phenotypes in adulthood. Here, we rear juveniles in either social groups or pairs to investigate the effects of early-life social environments on behavior and neuroendocrine gene expression. We find that both juvenile behavior and neuroendocrine function are sensitive to early-life effects. Behavior robustly co-varies across multiple contexts (open field, social cue investigation, and dominance behavior assays) to form a behavioral syndrome, with pair-reared juveniles towards the end of syndrome that is less active and socially interactive. Pair-reared juveniles also submit more readily as subordinates. In a separate cohort, we measured whole brain expression of stress and sex hormone genes. Expression of glucocorticoid receptor (GR) 1a was elevated in group-reared juveniles, supporting a highly-conserved role for the stress axis mediating early-life effects. The effect of rearing environment on androgen receptor (AR) α and estrogen receptor (ER) α expression was mediated by treatment duration (1 vs. 5 weeks). Finally, expression of corticotropin-releasing factor (CRF) and GR2 decreased significantly over time. Rearing environment also caused striking differences in gene co-expression, such that expression was tightly integrated in pair-reared juveniles, but not group-reared or isolates. Together, this research demonstrates the important developmental origins of behavioral phenotypes and identifies potential behavioral and neuroendocrine mechanisms.

## Introduction

Ontogeny has long been recognized as essential to understanding phenotype (Tinbergen, 1963), yet the early-life origins of individual behavioral variation remain understudied. Development reveals the proximate mechanisms by which genes interact with the environment during early life to sculpt the ‘machinery of behavior’ (Stamps, 2003; Tinbergen, 1963). Current or predicted environmental conditions can trigger developmental plasticity, and the resulting changes are often long-lasting, or even permanent, and can facilitate locally-adapted (e.g., predator resistant, Gilbert, 2001) phenotypes (Kasumovic and Brooks, 2011; Langenhof and Komdeur, 2018; Lummaa and Clutton-Brock, 2002; Piersma and Drent, 2003; Snell-Rood, 2013; Stamps, 2003; Stearns, 1989; West-Eberhard, 1989). The developmental mechanisms that shape social behavior via underlying neural regulatory mechanisms should be a particularly important target for natural selection (Taborsky, 2016) because of the direct consequences of social behavior for fitness and health (e.g., Bennett et al., 2006; Meyer-Lindenberg and Tost, 2012; Silk, 2007; Solomon-Lane et al., 2015; Wilson, 1980).

Social stimuli are among the most important attributes of the early-life environment (Taborsky, 2016). Although maternal (and, to a lesser extent, paternal) interactions have largely been the focus (e.g., Champagne & Curley, 2005; McClelland, Korosi, Cope, Ivy, & Baram, 2011), the broader early-life social environment is increasingly recognized for its role in behavioral and neural plasticity (Buist et al., 2013; Creel et al., 2013; Jonsson and Jonsson, 2014; Kasumovic and Brooks, 2011; Taborsky, 2016; White, 2010). For example, the early presence of brood care helpers, unrelated adult males, and multiple mothers and litters have long-term effects on social behavior in the Daffodil cichlid fish *Neolamprologus pulcher* (Arnold and Taborsky, 2010; Taborsky et al., 2012), brown-headed cowbirds (White et al., 2002), and laboratory mice (Branchi et al., 2013, 2006; D’Andrea et al., 2007), respectively. These features of the social environment alter the quality and quantity of social experiences and sensory cues perceived, which together influence neural function and behavior (Taborsky, 2016). Developmental plasticity may be limited to a single behavior or extend to an entire suite of behaviors (i.e., a behavioral syndrome), and the effects may be context-specific (Bell, 2007; Snell-Rood, 2013; Stamps, 2003; Stamps and Groothuis, 2010).

Neuroendocrine signaling is a primary mechanism by which environmental conditions and experience are translated into physiological responses (Crespi and Denver, 2005; Remage-Healey and Romero, 2000; Wingfield et al., 1990). Hormones are also important sources of individual variation in social behavior (e.g., across seasons, sexes, reproductive tactics) and underlie developmental plasticity relevant to adult behavior. The stress axis, or hypothalamic-pituitary-adrenal (interrenal in fish; HPA/I) axis, is widely implicated as a highly-conserved mechanism of early-life effects (Champagne and Curley, 2005; Francis et al., 1999; McClelland et al., 2011; Taborsky, 2016). In response to an environmental stressor, which includes any external condition that disrupts or threatens to disrupt homeostasis, the HPA/I axis integrates relevant internal and external cues and coordinates a response, such as changes in behavior and physiology. The stress response is initiated by the release of corticotropin-releasing factor (CRF) from the hypothalamus, which signals to the pituitary to release adrenocorticotropic hormone, which then signals the adrenal glands to release glucocorticoids (e.g., cortisol in fish) (Denver, 2009; Lowry and Moore, 2006; Wendelaar Bonga, 1997).

Effects of early-life experiences on HPA/I axis function have been demonstrated in every major vertebrate lineage (e.g., birds: Banerjee, Arterbery, Fergus, & Adkins-Regan, 2012; mammals: Champagne & Curley, 2005; amphibians: Crespi & Denver, 2005; fish: Jonsson & Jonsson, 2014). For example, the presence of brood helpers during early-life affects social behavior in the cooperatively breeding *N*. *pulcher* cichlid via changes in neural expression levels of CRF and glucocorticoid receptor (GR), as well as the ratio of the mineralocorticoid receptor (MR) to GR1 (Taborsky et al., 2013). Stress axis mechanisms can also mediate the effects of the early-life social environment on human health (e.g., Turecki & Meaney, 2016). Sex steroid hormones (e.g., androgens, estrogens) also play a role mediating the long-term effects of early-life experiences (Adkins-Regan, 2009; Brown and Spencer, 2013; Shepard et al., 2009) and regulating social behavior (Goodson, 2005; Newman, 1999). For example, neural estrogen receptor expression is associated with maternal behavior in mother rats and offspring (Cameron et al., 2008; Champagne et al., 2003; Champagne and Meaney, 2007), and socially stressed pre- and postnatal female guinea pigs have upregulated neural estrogen and androgen receptor levels, elevated testosterone, and masculinized behavior (Kaiser et al., 2003). Together, these and other neuroendocrine systems interact (e.g., Acevedo-Rodriguez et al., 2018) to affect behavior.

To investigate the effects of the early-life social environment on behavior and its neuroendocrine mechanisms, we used the highly social African cichlid *Astatotilapia burtoni*, a model system in social neuroscience (Fernald and Maruska, 2012; Hofmann, 2003; Stevenson et al., 2017). Adults of this species form mixed-sex, hierarchical communities with males of dominant or subordinate status and females. Dominant males are territorial, reproductively active, and colorful. In comparison, subordinate males shoal with females, are reproductively suppressed, and drab in coloration. Male status is socially regulated, and individuals regularly transition between status phenotypes (Fernald and Maruska, 2012; Hofmann, 2003). Adults, and juveniles (Fernald and Hirata, 1979), express a suite of highly evolutionarily conserved social behaviors, including aggression, affiliation, courtship, and cooperation (Fernald, 2012; Hofmann, 2003; Weitekamp et al., 2017). Substantial progress has also been made towards understanding variation in stress and sex steroid hormone signaling, including in the regulation of social behavior (Chen and Fernald, 2008; Fox et al., 1997; Greenwood et al., 2003; Munchrath and Hofmann, 2010; O’Connell and Hofmann, 2012a). All GRs (Greenwood et al., 2003), estrogen receptors (ER), and androgen receptors (AR) (Munchrath and Hofmann, 2010) have been studied in the adult *A*. *burtoni* brain, and neuroendocrine function can vary substantially. Subordinate males, for example, have lower levels of whole brain CRF and GR2 (Chen and Fernald, 2008), higher cortisol, and lower testosterone than dominants (Fox et al., 1997; O’Connell and Hofmann, 2012a), although these patterns can vary dynamically (Maguire and Hofmann, in prep.). The transcriptomic response in the preoptic area (POA) to pharmacological manipulation, such as an ER antagonist, is also status-specific (O’Connell and Hofmann, 2012a).

Given this rich literature on adult *A*. *burtoni*, it may seem surprising that the developmental origins of adult phenotypic variation remain largely unknown. The few studies that have investigated juveniles demonstrate the importance of early-life. For example, the development of male behavior and nuptial coloration, as well as reproductive maturation, are affected by the early-life social environment (Fernald and Hirata, 1979; Fraley and Fernald, 1982). Gestational cues (e.g., maternal social crowding) also have lasting effects on methylation and transcription of the *gnrh*1 gene in offspring (Alvarado et al., 2015). This result is particularly interesting given that POA GnRH1 neurons, which regulate gonadotropin release from the pituitary, are socially modulated in adults (Davis and Fernald, 1990; Hofmann and Fernald, 2001). However, studies of the effects of different early-life experiences on other neuroendocrine pathways or behavior are lacking.

In the present study, we conducted two experiments to test the hypothesis that the early-life social environment generates variation in juvenile behavior through neuroendocrine gene expression. We manipulated the early-life social environment, and consequently social experience, by rearing juveniles in either social groups or pairs. The natural distribution of territories and shoals across shallow shore pools and river estuaries (Fernald and Hirata, 1977; Rajkov et al., 2018) suggest that *A*. *burtoni* encounter a variety of dynamic social environments, including during development, although the degree of variation across individuals and over time has not been quantified. By directly manipulating group size, we can experimentally enhance the frequency, diversity, and/or complexity of early-life social experiences. Similar manipulations impact behavioral and neural development in a variety of species (reviewed in Taborsky, 2016). In the group condition, social experience implies interactions with more social partners, who also vary in size, sex, experience, and patterns of behavior. Interactions in groups can also involve more than two individuals, and it is possible to observe and learn from interactions of group members as a bystander. Although it has not been tested in juveniles, adults are capable of gaining important social information as a bystander (Desjardins et al., 2012, 2010; Grosenick et al., 2007). In the pair condition, juveniles occupy only one social role in a relationship with just one other individual. We predicted that rearing environment might affect social behavior— including social investigation, dominant, and subordinate behavior—possibly in a consistent manner across contexts. We also predicted effects on the expression of various genes that are part of candidate neuroendocrine systems known to mediate early-life experiences in other systems. Specifically, related to the HPA/I axis, we measured glucocorticoid receptor 1a (GR1a), glucocorticoid receptor 1b (GR1b), glucocorticoid receptor 2 (GR2) (nomenclature from Maruska & Fernald, 2010), mineralocorticoid receptor (MR), and CRF. For sex steroid hormone signaling, we quantified androgen receptor α (ARα) and estrogen receptor α (ERα). By investigating these early-life effects in juveniles, we can identify important intermediary steps that inform how developmental plasticity may shape the adult phenotype.

## Methods

### Animals

Juvenile *A*. *burtoni* came from a laboratory population descended from a wild-caught stock. The adults that bred the juveniles were housed in naturalistic social groups of males and females. Dominant males court gravid females that then lay eggs in his territory. The female then scoops up the eggs into her mouth, where the male fertilizes them. The mother orally incubates the larvae as they develop for 10-13 days. Under natural (and some laboratory) conditions, juveniles remain close to their mother for the 2-4 weeks following their initial release from her mouth. As they age, juveniles seek shelter in her mouth less and less often. In the first two weeks, juveniles primarily school together, with overt social interactions beginning at 2-3 weeks old (Fernald and Hirata, 1979; Renn et al., 2009). Social behaviors, such as chasing, nipping, territorial displays, emerge in a predictable sequence as juveniles approach reproductive maturity, which can occur as early as 15 weeks, depending on the early-life social conditions (Fernald and Hirata, 1979; Fraley and Fernald, 1982).

We removed juveniles from the mother’s mouth 6-12 days post-fertilization. Once sufficiently developed (∼day 12, freely swimming with no remaining yolk), juveniles were transferred into experimental rearing environments. Juveniles are all silver (drab) in coloration, and none developed coloration during the study, which would indicate reproductive maturity for males. Sex cannot be determined anatomically until maturation; therefore, the sex ratios of our rearing environments, and the sex of the focal individuals, is unknown. The sex ratio of *A*.*burtoni* broods is approximately 1:1 (Heule et al., 2014). All work was done in compliance with the Institutional Animal Care and Use Committee at The University of Texas at Austin.

### Experimental rearing conditions (Experiments 1 & 2)

As the first study of this kind in this species, we opted to quantify behavior and gene expression in separate experiments in order to capture different developmental time points. In Experiment 1, juveniles for the behavioral assays were reared in social groups of 16 fish (n=12 groups) or in pairs (n=9 pairs) for 58-73 days (average 65.76 ± 0.81; ∼8-10 weeks). This is the longest duration that could be used without juveniles reaching reproductive maturity. In Experiment 2, neural gene expression was measured in a separate cohort of juveniles reared in social groups of 16 fish, pairs, or in isolation for 1 week (groups: n=8; pairs: n=8; isolates: n=8) or 5 weeks (groups: n=14; pairs: n=10). Here, we aimed to capture early changes in gene expression that might set individuals along different developmental trajectories. Isolation was included because we expected it to impact gene expression in this highly social species, not as a social control. We cannot distinguish between the effects of chronological age from the treatment duration (1 vs. 5 weeks) in this study.

For both Experiments, juveniles from multiple clutches of the same age and developmental stage (day 12-14 fry) were divided among treatment groups. Group-reared fish were housed in 35 L aquaria with three terracotta pot shards for shelter and/or territory. Pairs and isolated fish were housed in small aquaria (22.9 x 15.2 x 15.2 cm) with one terracotta pot shard. The volume of water per fish was similar for the group (2.6 L) and paired (2.7 L) treatments. Juveniles were fed daily with Hikari plankton (Pentair Aquatic Eco-Systems, Cary, NC). The food was mixed in water, and a transfer pipette was used to deliver a set volume to each tank.

Groups received eight times more food than pairs. Pairs and isolated fish received the same amount. All juveniles were maintained on a 12:12 light/dark cycle.

### Experiment 1: Behavioral assays

We quantified behavior in four assays, which were always presented in the same sequence (Fig 1): an open field test that is commonly used in other species to assess activity and anxiety (e.g., Cachat et al., 2010; Prut & Belzung, 2003); a social cue investigation as a measure of social motivation or preference (e.g., Bonuti & Morato, 2018; Moy et al., 2004); and social interactions within either dominant or subordinate status contexts, which individuals regularly experience in social communities of *A*. *burtoni* (Hofmann, 2003). Behavioral neuroscientists employ a wide range of different assays across different model systems, and we explored which assays juvenile *A*. *burtoni* would participate in in a series of pilot experiments. We decided on this combination of assays because each assay has been used with multiple species, thus allowing for cross-species comparisons, and the target behaviors (e.g., locomotion, space use, social approach, social interaction) are all expressed by *A*. *burtoni* in natural contexts and directly relevant to adult social status and reproduction (e.g., via territoriality, aggression). Including multiple assays in combination also provides a more comprehensive understanding of behavioral phenotype, which is complex and expressed in context-specific ways.

**Figure 1:**
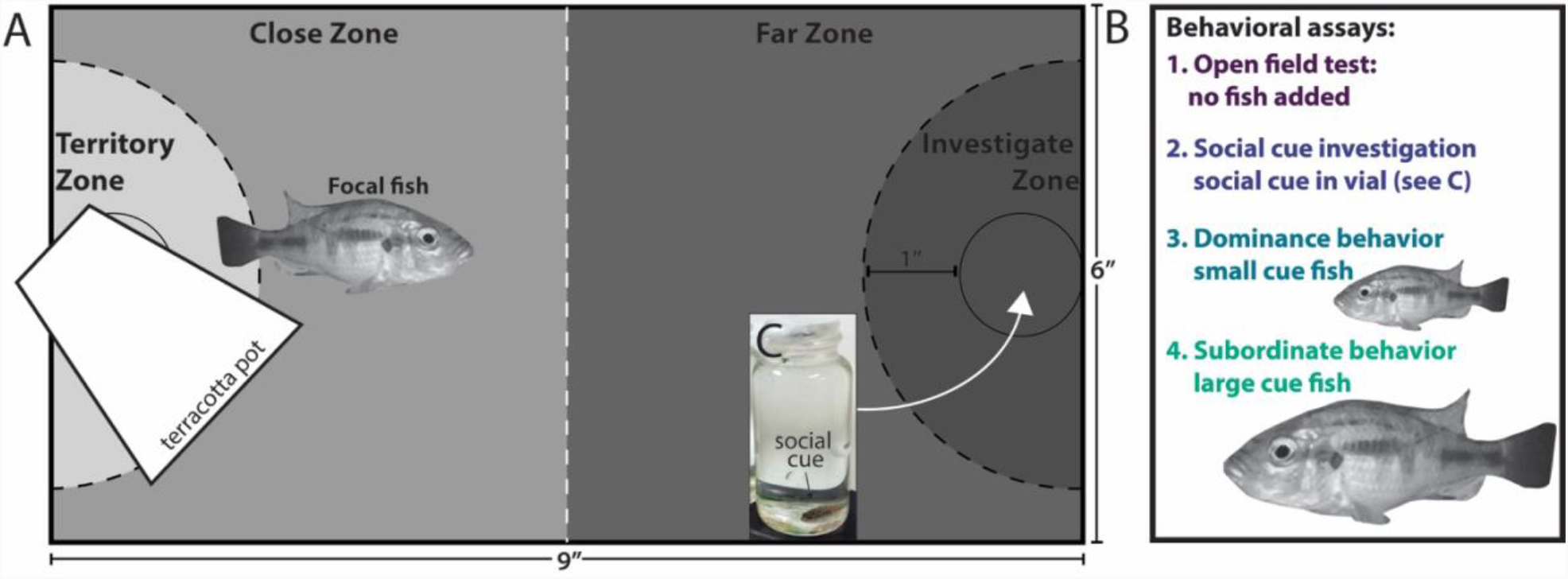
Experimental setup for behavior assays. Juvenile behavior was observed in a novel experimental tank in four sequential assays administered in the same order, each lasting 30 min. A terracotta shard served as a shelter and/or territory. The black lines (dotted, solid) were drawn on the tank bottom in permanent maker, dividing the tank into four zones: territory, close, far, and investigate. The center dividing line (white) was not drawn (A). The focal fish was alone in the tank for the open field assay, and the time in each zone and frequency of entered each zone was recorded (B, assay 1). For the social cue investigation, a juvenile inside of a scintillation vial was placed in the circle within the investigate zone (see C). The time in and frequency of entering each zone was recorded (B, assay 2). The social cue was removed and a freely swimming, novel cue fish (smaller than the focal) was added to the tank for the dominance behavior assay (B, assay 3). The small cue fish was then removed and a freely swimming, novel cue fish (larger than the focal) was added to the tank for the subordinate behavior assay (B, assay 4). Social interactions were recorded for the dominant and subordinate behavior assays. The time in and frequency of entering the territory zone was also recorded for both fish.

Behavior for both members of the pairs (n=18 individuals) and two fish from each group (n=24 individuals) was analyzed. To choose focal individuals from the groups, we removed all fish from the aquarium and selected, by eye, one of the largest fish. A smaller fish was then chosen such that the ratio of large-to-small fish standard length (SL, mm) was approximately equal in the group and a pair from the same cohort of juveniles (same age). These smaller fish were never the smallest in their groups. Because size is a strong predicator of social status (Alcazar et al., 2014), the larger fish was very likely to have dominance experience, similar to the larger fish in the pair. The smaller fish were very likely to have subordinate and dominant interactions with larger and smaller individuals in the group, respectively. Standard length was recorded for all focal fish.

Behavior was observed in novel, small aquaria (22.9 x 15.2 x 15.2 cm) without covers. For analysis, the aquaria were divided into 4 zones (Fig 1), delineated with permanent marker. In the middle of each short side, a circle was drawn (28 mm diameter) to indicate the placement of the scintillation vial (see below: social cue investigation). An arc 2.54 cm from the edge of that circle was drawn to form a semicircle. One semicircle was designated the “territory” zone and had a terracotta pot shard for a shelter and/or territory. The other semicircle was designated the “investigate” zone. The “close” zone was between the territory zone and halfway along the long side of the tank. The “far” zone was between the halfway mark and the investigate zone (Fig 1). Video cameras recorded behavior from above so that all areas of the tank, except under the terracotta pot shard, were visible. Solomon Coder was used for analysis (www.solomoncoder.com). All observations were made by the same observer who was blind to treatment. Ten minutes of behavior was analyzed from each behavior assay for a total of 40 min of behavior scored for each individual.

#### Open field test

The focal fish was transferred to the test aquarium with a hand net and remained in the tank alone for 30 min. Movement around the tank was observed from minutes 20 to 30. We recorded the number of times a fish crossed into each zone (frequency) and the time (s) spent in each zone. <u>Social cue investigation</u>: Novel juveniles were collected from a community tank and placed into scintillation vials (20 mL). The top of the vial was covered with parafilm with holes to allow water through. A vial containing one cue fish was placed into each test aquarium (n=16 group-reared, n=13 pair-reared). Cue fish were 0-6.4 mm SL (average 3.37 ± 0.27) smaller than their focal fish. An empty vial was used as a control (n=8 group-reared, n=5 pair-reared). The social cues were in the aquarium for 30 min. Movement around the tank (frequency and time in each zone) was scored from minutes 2 to 12.

#### Dominance behavior

The scintillation vials were removed from the aquaria and a novel smaller fish (by 1-6.4 mm SL, average 3.37 ± 0.25) was immediately added to each aquarium, freely swimming with the focal fish. The pair remained together for 30 minutes, and behavior was scored from minutes 2 to 12. <u>Subordinate behavior</u>: The small cue fish was removed from the aquaria and a novel, larger fish (by 2.4-12 mm SL, average 5.74 ± 0.34) was immediately added to each aquarium, freely swimming with the focal fish. The pair remained together for 30 minutes, and behavior was scored from minutes 2 to 12. In the dominance and subordinate behavior assays, we analyzed agonistic interactions between the pair. An approach was defined as one fish swimming directly towards any part of the other fish’s body, within 3 body lengths. If the approached fish responded by moving away, in any direction, the behavior was recorded as a displacement for the initiator and a submission for the responder. From these measures, we calculated agonistic efficiency, or the proportion of approaches that led to a displacement (Solomon-Lane et al., 2014), for focal and cue fish. The difference in agonistic efficiency between the focal and cue fish was used as a measure of agonistic asymmetry, which characterizes status relationships (Drews, 1993). We also recorded the frequency of entering and the time spent in the territory, for the focal fish, cue fish, and both together.

### Experiment 2: Whole brain gene expression

Whole brain gene expression for two fish from each group (1 week: n=8; 5 weeks: n=14), both members of the pairs (1 week: n=8; 5 weeks: n=10), and every isolate (1 week: n=8) was analyzed. Because the present study is the first to examine the neuromolecular substrates associated with early life social experience in *A*. *burtoni*, we did not have an *a priori* expectation as to which brain regions or cell types might be the most critical to examine. Therefore, we decided to analyze expression in whole brain, even though important differences in circuits and brain regions may not be identified using this approach. It should also be noted that recent evidence suggests that patterns of expression specific to brain region, or even cell-type, can be inferred from bulk tissue samples (Kelley et al., 2018), such as whole brain.

Focal individuals from the group condition were selected haphazardly. Juveniles were removed from their rearing environments with a hand net and rapidly decapitated. The brains were dissected immediately, flash frozen on dry ice, and stored at −80°C until processing. Gene expression was quantified using qPCR and previously validated primers (Supplemental Table 1, Chen and Fernald, 2008; Greenwood et al., 2003; O’Connell and Hofmann, 2012a) for GR1, GR2a, GR2b, MR, CRF, ARα, and ERα, as well as control genes 18S and G3PDH. With regards to (nuclear) sex steroid receptors, we chose subtypes ARα and ERα because of their demonstrated role in the regulation of adult *A*. *burtoni* social and reproductive behavior (Burmeister et al., 2007; Korzan et al., 2014; Maruska, 2015; O’Connell and Hofmann, 2012a). Other subtypes (e.g., ARβ, ERβ a,b), as well as progesterone receptor, also have distinct distributions and regulatory roles (Burmeister et al., 2007; Munchrath and Hofmann, 2010) and certainly warrant investigation in future studies. RNA was extracted using the Maxwell 16 LEV simplyRNA Tissue Kit (Promega, Madison, WI), and the Promega GoScript Reverse Transcription System (Promega, Madison, WI) was used for reverse transcription. PowerUp SYBR Green Master Mix (ThermoFisher Scientific, Waltham, MA) was used for quantitative PCR. All standard kit protocols were followed. Relative gene expression levels were quantified using ΔΔCT analysis, using 18S and G3PDH as reference genes. The results are largely concordant independent of the reference gene used. Here, we present the analyses for 18S, as this gene has shown very little expression variation across social phenotypes in transcriptome studies of *A*. *burtoni* (O’Connell and Hofmann, 2012a; Renn et al., 2008).

### Statistical analyses

All statistical analyses were conducted using R Studio (version 1.0.143). Results were considered significant at the p<0.05 level, and averages ± standard error of the mean are included in the text. Cohen’s d is reported to estimate effect size (small effect: 0.2<d<0.5; medium: 0.5<d<0.8; large: 0.8<d). The box of the box and whisker plots show the median and the first and third quartiles. The whiskers extend to the largest and smallest observations within or equal to 1.5 times the interquartile range. Comparisons between group- and pair-reared juveniles were conducted using t-tests for fish SL, time and frequency in each tank zone, and rates of agonistic behavior. Mann-Whitney-Wilcoxon tests were used for data that did not meet the assumptions of parametric statistics. Regression analysis was used to identify significant associations between SL and frequency and time in a zone and between SL and agonistic behavior. We used a false discovery rate correction for regressions with focal fish SL (Benjamini and Hochberg, 1995). Two-way ANOVAs were used to identify significant effects of rearing environment, presence of the social cue, or an interaction, on the frequency and time spent in each zone of the tank. We used Principal Components Analysis (PCA) to identify how behaviors clustered across the four assays and for each assay individually. Independent t-tests were used to compare principal component scores between group- and pair reared juveniles. Paired t-tests (or Mann-Whitney-Wilcoxon tests) were used to compare principal component scores between the larger and smaller fish sampled from groups and pairs. Correlation analysis was used to identify significant associations among principal components (PCs).

We used two-way ANOVAs to identify significant effects of rearing environment (group, pair, isolated), treatment duration (1 week, 5 weeks), or an interaction on the expression of individual candidate genes. All gene expression data were log transformed to meet the assumptions of parametric statistics. Partial correlation networks were calculated using the “ppcor” package in R and visualized using “qgraph.” The nodes of the networks represent the gene. The edges are the partial correlation coefficient, with thicker edges indicating stronger correlations. Only significant correlations are shown. Mantel tests were used to test for pairwise differences between the gene expression networks. A non-significant p-value (> 0.05) indicates that the partial correlation matrices are not related.

## Results

### Experiment 1

#### Standard length

After 8-10 weeks in their respective treatment condition, group-reared juveniles (16.85 ± 0.32 mm SL) were significantly larger than pair-reared juveniles (13.76 ± 0.40 mm SL) (t=6.00, p=7.25e-7, d=1.89). This size difference influenced the size of the fish selected to be the social stimuli. Specifically, the difference in SL between the focal fish and the social cue (t=3.38, p=0.0016, d=1.02), as well as the focal fish and the small cue fish (t=3.48, p=0.0013, d=1.09), was significantly greater for group-reared juveniles. The SL difference between the focal fish and the large cue fish was significantly greater for pair-reared juveniles (t=-3.22, p=0.0025, d=0.95). Relative size differences followed the same pattern as absolute size differences (data not shown).

#### Open field test and social cue investigation

In the open field test (and subsequent assays), juveniles of both treatment groups moved readily around the novel environment with minimal acclimation. We present the data for the frequency of entering each zone (Supplemental Fig 1A-D). There were no significant effects for the time spent in each zone (p>0.05). Group-reared juveniles entered the territory (Mann-Whitney-Wilcoxon test: W=299, p=0.034, d=0.51), close (W=293.5, p=0.049, d=0.41), and investigate zones (W=293.5, p=0.049, d=0.60) significantly more frequently than pair-reared juveniles. There was no significant difference for the far zone (W=289, p=0.064).

Next, we used a social cue investigation task to examine whether and how rearing environment and/or the presence of the social cue affect locomotor activity (Supplemental Fig 1E-H). Two-way ANOVA revealed that, following the addition of the social cue, juveniles entered the investigate zone significantly more frequently than controls (F_1,36_= 4.91, p=0.033, d=0.96). There was no effect of rearing environment (F_1,36_ =1.69, p=0.20) and no interaction (F_1,36_=0.046, p=0.83). There was no effect of rearing environment (F_1,36_ =2.68, p=0.11), social cue (F_1,36_ =0.87, p=0.36), or an interaction (F1,36=0.84, p=0.37) on frequency of entering the far zone. Group-reared juveniles entered the close zone significantly more than pair-reared juveniles (F_1,35_=4.47, p=0.042, d=0.71), but there was no effect of the social cue (F_1,35_=0.11, p=0.74) and no interaction (F_1,35_=0.44, p=0.52). There was no effect of rearing environment (F_1,35_=3.28, p=0.079), social cue (F_1,35_=0.17, p=0.68) and no interaction (F_1,35_=0.83, p=0.37) on the frequency of entering the territory zone. Linear regression analyses show that SL is not associated with the frequency of entering zones of the tank for group-or pair-reared juveniles (Supplemental Table 2).

#### Dominant and subordinate behavior

Rearing environment did not affect rates of focal fish behavior (Supplemental Fig 2). As the dominant fish, there were no differences in approaching (W=242.5, p=0.20) or displacing (W=253, p=0.12) the small cue fish. As the subordinate, there were no differences in approaching (W=205.5, p=0.85), displacing (W=214.5, p=0.62), or submitting to (W=217.5, p=0.56) the large cue fish. In the dominance assay, rearing environment did not affect agonistic efficiency for the focal fish (t=0.83, p=0.41), small cue fish (W=115.5, p= 0.97), or the difference between the pair (t=1.03, p=0.32). In the subordinate assay, although there was no effect of rearing environment on agonistic efficiency for the focal fish (W=169.5, p=0.28) or the large cue fish (W=112.5, p=0.061), the difference in agonistic efficiency was significantly higher for pair-reared juveniles (t=-2.42, p=0.022, d=0.81). Linear regression analyses show that SL is not associated with social behavior for group- or pair-reared juveniles (Supplemental Table 2).

#### Multivariate analysis of behavior across assays

In order to gain more insight into this multivariate dataset, we employed PCA to determine which measures of morphology (i.e., size) and behavior might act in concert to explain different aspects of the variability across individuals, including based on rearing environment and whether the focal individual was the larger or smaller fish sampled from the group or pair.

Given that body size serves as a reliable proxy for social status experience in adults, we refer to the larger and smaller juvenile as dominant and subordinate, respectively. We first conducted a PCA that included variables from each of the four assays: focal fish SL; frequency of entering each zone in the open field test and social cue investigation; focal fish social approaches and displacements as a dominant towards the small cue fish; and focal fish approaches, displacements, and submissions as a subordinate with the larger cue fish. We found that principal component (PC) 1 accounts for 43.3% of the total variance and differs significantly between group- and pair-reared juveniles (t=-2.30, p=0.029, d=0.75, Fig 2A). There was a trend for PC2 (16.4%; z= −1.96, p=0.05, d=0.39, Fig 2B) to differ based on status experience (or relative size), and the difference was significant for PC5 (6.6%; t=--2.16, p=0.043, d=0.53, Fig 2C). PC6 (5.0%) also differed significantly between group- and pair-reared juveniles (t=4.66, p= 4.082e-5, d=1.46, Fig 2D). No significant differences were identified for other PCs (p>0.05). As the vector plot in Fig 2E shows, variables from the open field test, social cue investigation, and dominance behavior assay all load on PC1, along with focal fish SL, while measures of behavior during the subordinate assay load on PC2. The vector plot in Fig 2F shows that a number of behaviors load on PC5, the strongest of which relate movement around the tank during the open field and social cue investigation assays. Focal fish SL loads most strongly on PC6.

**Figure 2:**
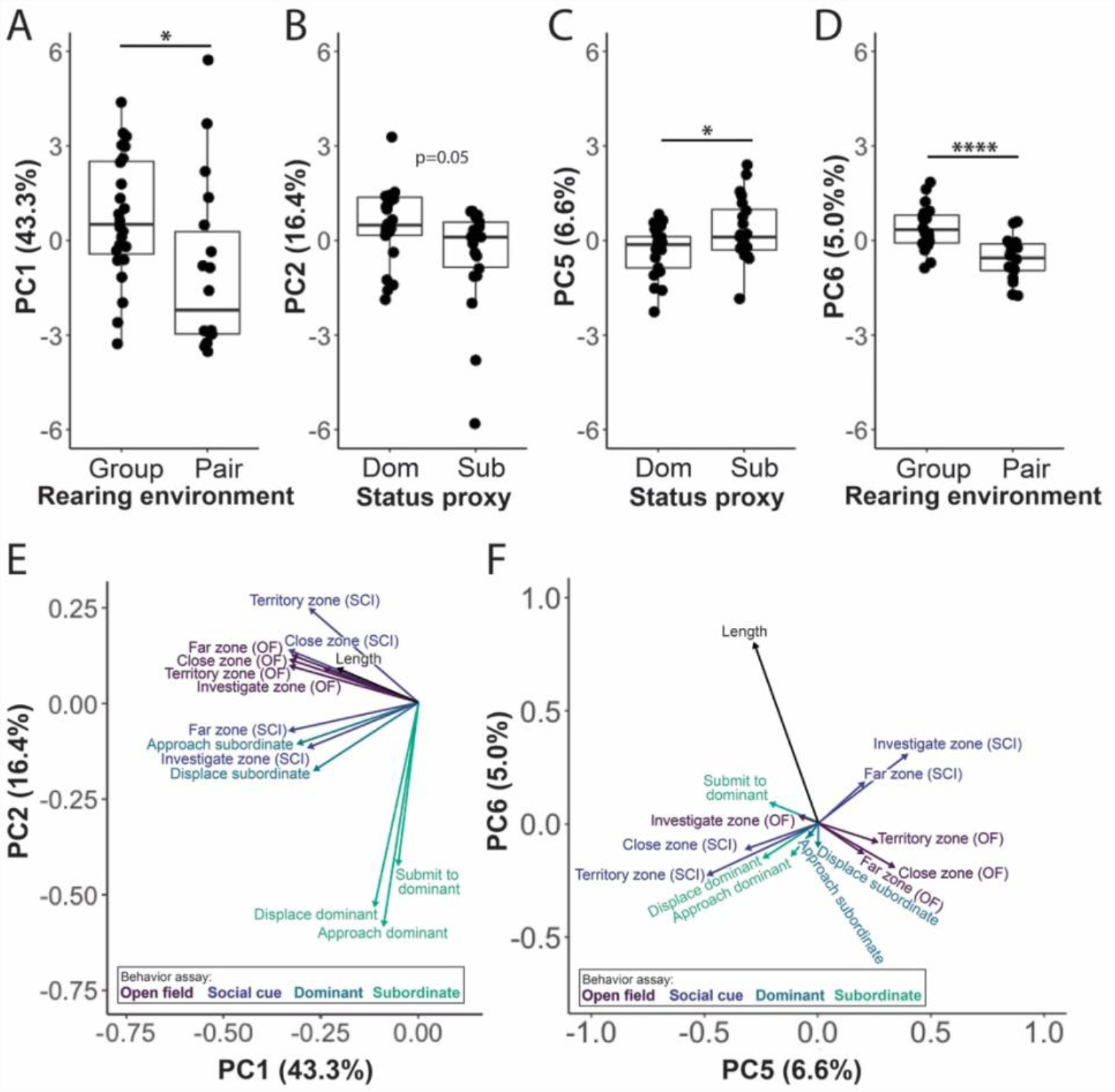
Principal component analysis (PCA) of focal fish behavior from all four assays (open field, social cue investigation, dominance, subordinate behavior). Differences in PC1 between group- and pair-reared juveniles (A). Differences in PC2 (B) and PC5 (C) between the larger / dominant (Dom) fish and smaller / subordinate (Sub) fish selected from the group and pair. The larger fish is very likely to have more dominance experience, while the smaller fish has more subordinate experience. Differences in PC6 between group- and pair-reared juveniles (D). Vector plot showing the PCA variables that load on PC1 and PC2 (E). Vector plot showing the PCA variables that load on PC5 and PC6 (F). Percentages refer to the amount of variance explained by that component. Pair (n=18 individuals). Group (n=24 individuals). Social cue investigation (SCI). Open field exploration (OF). *p<0.05, ****p<0.001.

To disentangle the possible effects of SL and rearing environment on behavior, we re-ran the PCA without focal fish SL. In this analysis, PC1 (44.8% of the variance) still differs significantly between group-reared and pair-reared juveniles (W=126, p=0.022, d=0.64). Although focal fish SL is significantly and positively correlated with PC1 (r^2^=0.19, p=0.0026), SL does not correlate with PC1 for group-reared (p=0.16) or pair-reared juveniles (p=0.096) separately.

To better understand how rearing environment affected behavior within the assays that contributed to the treatment difference in PC1 (Fig 2E), we conducted PCAs for the open field, social cue investigation, and dominance behavior assays separately. We expanded these analyses to include all of the measured variables, for the focal and cue fish. The open field test analysis included focal fish SL and the frequency of entering and time in each zone of the tank. The social cue investigation included the same measures, as well as the SL of the cue fish. Finally, the dominance behavior analysis included SL of the focal fish and small cue fish, approaches and displacements of both fish, and the frequency of entering and time spent in the territory by either or both fish. For each analysis, we focused on PC1, which differed significantly between group- and pair-reared juveniles: open field (accounting for 43.4% of the total variance; t=-2.14, p=0.04, d=0.71, Fig 3A), social cue investigation (37.2%; W=102, p = 0.0032, d=0.92, Fig 3B), and dominance behavior (29.8%; W=128, p=0.025, d=0.71, Fig 3C). The PC1s were also significantly and linearly correlated with each other (Fig 3D, open field x social cue: r^2^=0.46, p=5.33e-7; open field x dominance: r^2^=0.33, p= 4.69e-5; social cue x dominance: r^2^=0.46, p= 4.97e-7, Supplemental Fig 3). See Supplemental Figure 4 for the proportion of the variance explained by each PC. Vector plots in Supplemental Figure 5 shows the variables that load on PC1 (and PC2) for each included assay.

**Figure 3:**
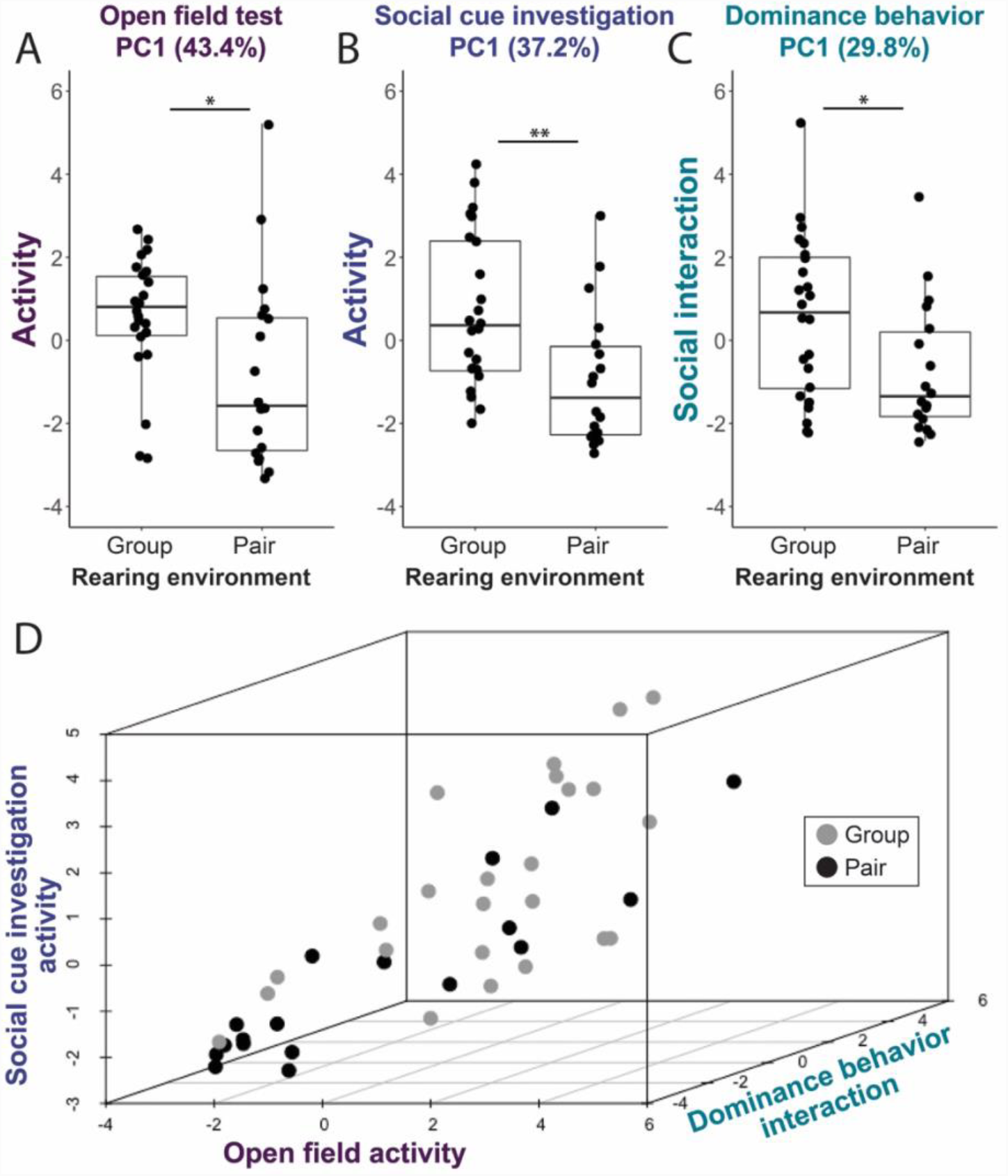
Separate principal component analyses performed for the open field (A), social cue investigation (B), and dominance behavior (C) assays. Both focal and non-focal fish variables (behavior, size). The significant, positive correlations about the PC1s are shown in a three-dimensional plot (D). Percentages refer to the amount of variance explained by that component. Pair (n=18 individuals). Group (n=24 individuals). *p<0.05, **p<0.01.

### Experiment 2

#### Neural gene expression patterns

Neuroendocrine signaling is a primary mechanism by which early-life experiences are translated into biological changes. To identify potential mediators of the behavioral effects we identified, we measured mRNA levels of genes involved in the stress axis and in sex steroid signaling in the brains of a separate cohort of juveniles. We compared relative expression across rearing environments (isolation, pairs, groups) and time in rearing environment (1 week, 5 weeks) (Fig 4) using two-way ANOVAs. The sex steroid hormones, ARα and ERα, were the only genes to have significant interactions between rearing environment and treatment duration. For ARα, there was no significant effect of treatment (F_2,42_=2.23, p=0.12), but there was a significant effect of treatment duration (F1,42=7.89, p=0.0075) and a significant interaction (F_1,42_=4.95, p=0.032). *Post hoc* analysis of the simple main effects revealed that for the 5 week juveniles, ARα expression was significantly higher in group-reared fish (t=3.67, p=0.0015).

**Figure 4:**
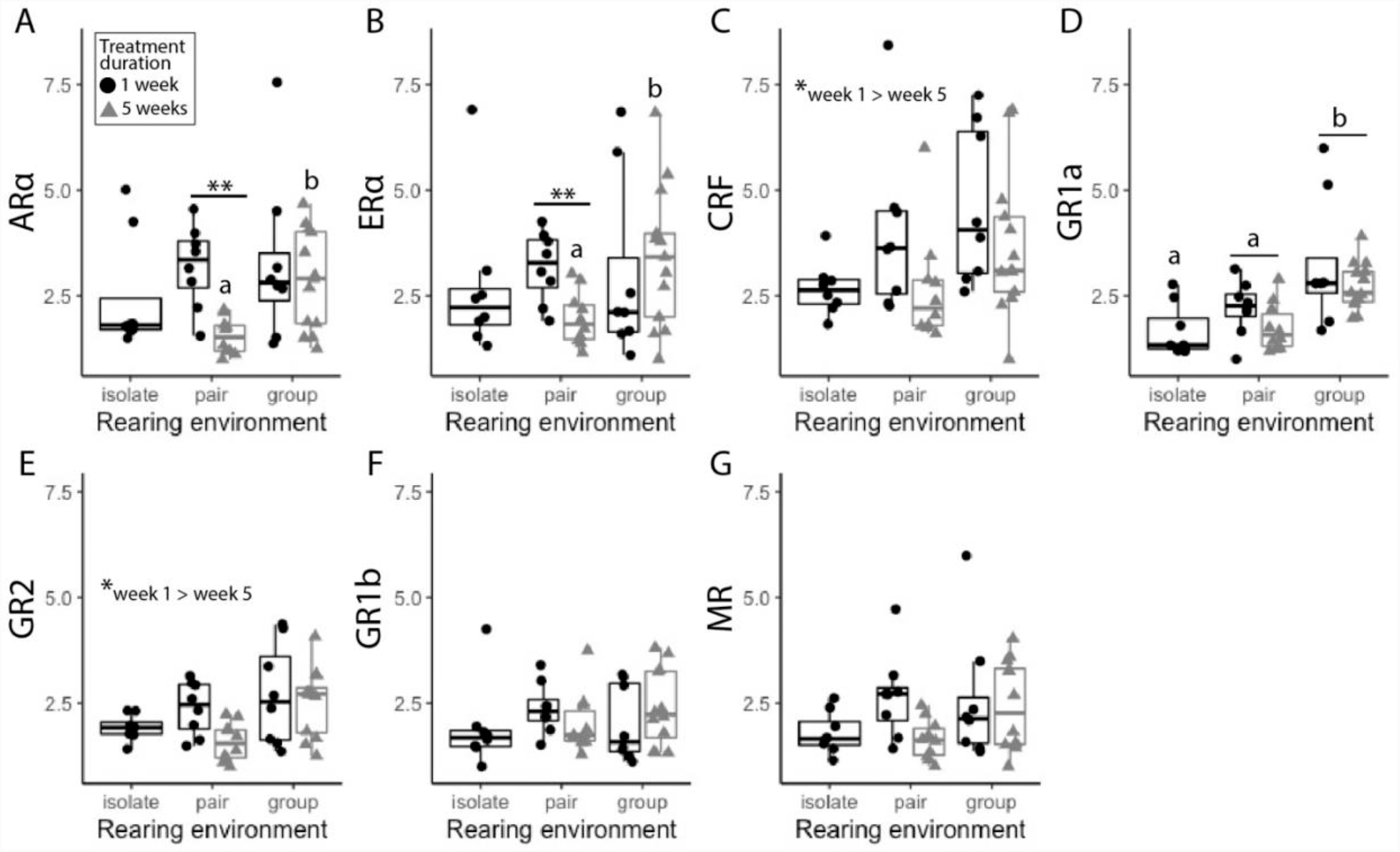
Relative gene expression calculated using ΔΔCT analysis (reference gene 18S) for juveniles reared in isolation (1 week, n=8), pairs (1 week or 5 weeks, n=18), and groups (1 week or 5 weeks, n=22). Androgen receptor α (ARα). Estrogen receptor α (ERα). Glucocorticoid receptors (GR). Mineralocorticoid receptor (MR). Corticotropin-releasing factor (CRF). Letters indicate significant differences across treatment groups (p<0.05). *p<0.05, **p<0.01.

There were no treatment differences after 1 week (F_2,21_=1.15, p=0.34). In pair-reared juveniles, ARα expression was significantly higher after 1 week in treatment compared to after 5 weeks (t=4.72, p=0.00038). There were no treatment duration differences among group-reared juveniles (t=0.42, p=0.68), and isolates were only analyzed following 1 week in treatment, so comparison was not possible (Fig 4A). For ERα, there was no significant effect of treatment (F_2,42_=0.73, p=0.49) or treatment duration (F_1,42_=0.71, p=0.41), but there was a significant interaction (F1,42=4.89, p=0.032). *Post hoc* analysis of the simple main effects revealed a pattern similar to ARα. For juveniles in treatment groups for 5 weeks, ERα expression was significantly higher for group-reared juveniles (t=2.59, p=0.018). There were no differences after 1 week in treatment groups (F_2,21_=0.63, p=0.54). In pair reared juveniles, ERα was significantly higher after 1 week in treatment compared to after 5 weeks (t=3.49, p=0.0031). There were no treatment differences among group-reared juveniles (t=-0.73, p=0.48) (Fig 4B).

For genes related to the stress response, we found significant main effects for CRF, GR1a, and GR2. For CRF, there was a significant effect of treatment duration, where week 1 expression was significantly higher than after 5 weeks in treatment F_1,42_=5.77, p=0.021). There was no effect of treatment (F_2,42_=2.45, p=0.099) and no interaction effect (F_1,42_=0.27, p=0.61) (Fig 4C). For GR1a, there was a significant effect of treatment (F_2,42_=12.47, p=5.63e-5), and *post hoc* analysis showed that group-reared juveniles had significantly higher expression than pair-reared (p=0.0008) and isolated (p=0.00034) juveniles. Expression for pair-reared juveniles was not significantly different from isolates (p=0.49). There was no main effect of treatment duration (F_1,42_=2.32, p=0.14), and there was no interaction (F_1,42_=0.38, p=0.54) (Fig 4D). For GR2, there was a significant main effect of treatment duration (F_1,42_=4.10, p=0.049), and similar to CRF, expression was significantly higher after 1 week in treatment. There was also a significant main effect of treatment (F_2,42_=3.40, p=0.026); however, *post hoc* analysis revealed that none of the pairwise differences were significant (group vs. isolates: p=0.20; group vs. pair: p=0.084; pair vs. isolate: p=0.85). The interaction effect was not significant (F_1,42_=3.25, p=0.079) (Fig 4F). There were no significant differences for GR1b (treatment: F_2,42_=0.70, p=0.50; treatment duration: F_1,42_=0.01, p=0.92; interaction: F_1,42_=2.38, p=0.13; Fig 4E) or MR (treatment: F_2,42_=1.32, p=0.28; treatment duration: F_1,42_=3.28, p=0.077; interaction: F_1,42_=2.91, p=0.095; Fig 4G).

Genes function within regulatory networks, rather than in isolation, and they can affect each other’s expression. A common upstream regulator may also control multiple functional networks of genes. Because of their known effects on physiology and behavior, these candidate genes are likely to function in pathways that interact with each other. To quantify how rearing environment affects gene co-expression, we calculated partial correlation networks (Fig 5). Partial correlations show the associations between gene pairs, independent of other correlations in the network. Comparing the group and pair networks (Mantel test: p=0.31), the group and isolate networks (p=0.61), and the pair and isolate networks (p=0.12) revealed that there was no evidence that any of these networks were similar to any other.

**Figure 5:**
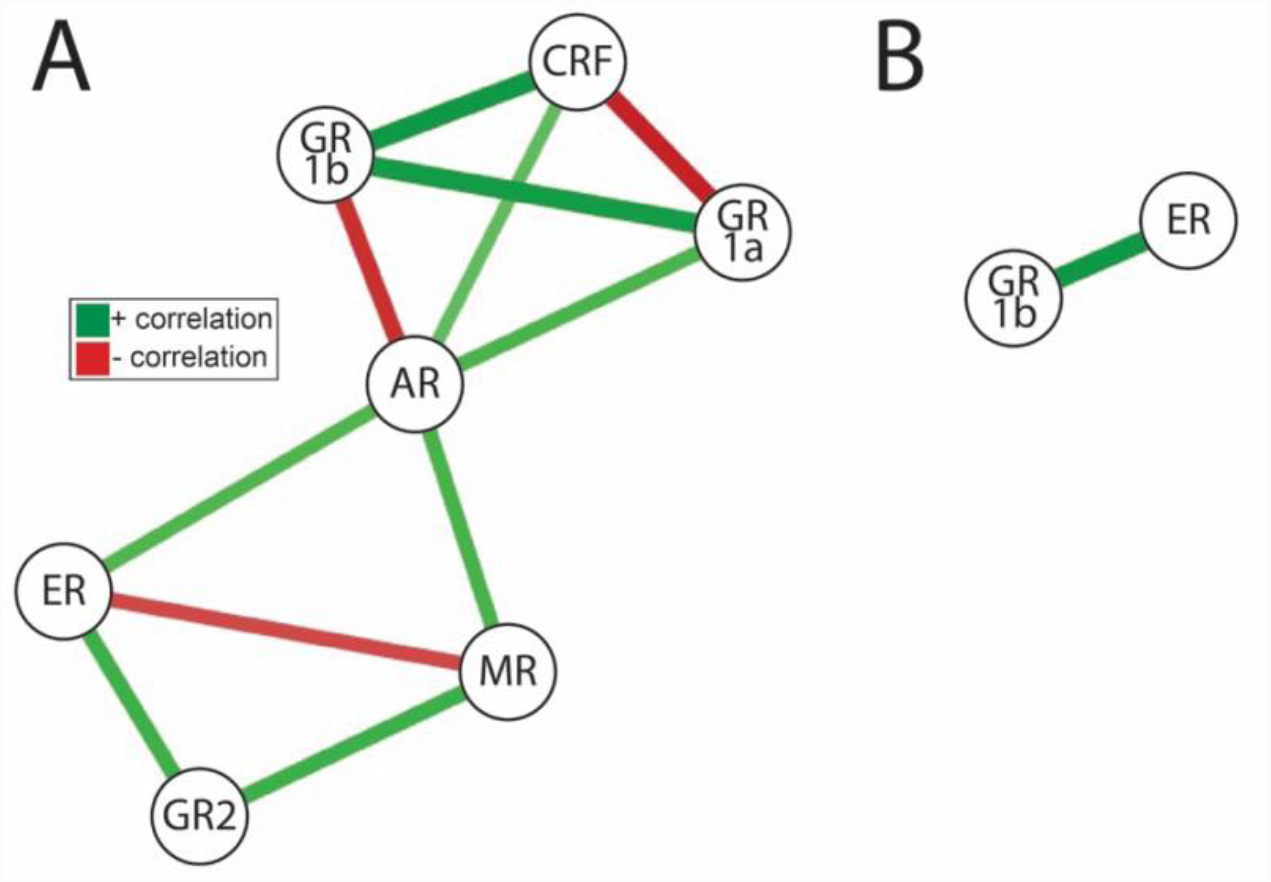
Partial correlation network of gene expression in pair-reared juveniles (n=18) (A) and group-reared juveniles (n=22) (B). Nodes are the candidate genes. Edges represent partial correlations between nodes. Only significant partial correlations are shown (p<0.05), and edge thickness indicates correlation strength. There were no significant partial correlations for juveniles reared in isolation (n=8) (p>0.05). Androgen receptor α (AR). Estrogen receptor α (ER). Glucocorticoid receptors (GR). Mineralocorticoid receptor (MR). Corticotropin-releasing factor (CRF).

## Discussion

In the present study, we demonstrate that juvenile *A*. *burtoni* behavior and neuroendocrine gene expression are both sensitive to early-life social effects. By rearing juveniles in different social environments—either in a social group or as a pair, both of which allow individuals to interact freely at all times—we altered the quality and quantity of social experiences and sensory cues perceived and set individuals along different developmental trajectories. Behaviorally, the early-life environment shifted juveniles in a predictable manner along a continuum of a novel behavioral syndrome (i.e., correlated behaviors across contexts, see below) comprised of open field, social cue investigation, and dominance behaviors (Fig 2, Fig 3) and affected patterns of subordinate behavior, a critically important social role for young individuals. In the brain, rearing environment caused significant changes in the expression of key neuroendocrine genes, including ARα, ERα, and GR1a (Fig 4), and led to striking differences in patterns of co-expression (Fig 5). The significant effects of treatment duration also provide important insights into developmental processes (Fig 4). Together, these experiments provide an essential step towards understanding how developmental plasticity generates the individual variation in behavior and neuroendocrine function that has fitness and health consequences in adulthood (e.g., Champagne, 2010; Turecki and Meaney, 2016). Our results also contribute to an important and growing literature on the impact of early-life social environments beyond parental interactions (Champagne and Curley, 2005; Taborsky, 2016), using a species that, despite its prominence in social neuroscience (Fernald and Maruska, 2012; Hofmann, 2003), has rarely been studied during development (Alvarado, Lenkov, Williams, & Fernald, 2015; Fernald & Hirata, 1979; Fraley & Fernald, 1982).

### Juvenile behavior forms a syndrome affected by early-life social environment

Using a battery of four behavioral assays to gain a comprehensive understanding of behavioral phenotype, within and across contexts (Fig 1), we discovered that open field, social cue investigation, and dominance behavior together formed a behavioral syndrome (Fig 3). Syndromes are a population-level metric defined as the correlation between rank-order differences between individuals, across contexts and/or over time (Bell, 2007). The presence of a syndrome indicates consistency in patterns of individual behavior across contexts and/or over time (Bell, 2007; Sih et al., 2004b, 2004a). Our data suggest that how individuals move around in space is relevant to the social role they play. Specifically, juveniles that were more active in the open field test were more likely to be active in the social cue investigation and more interactive in the dominance assay (Fig 3). Interestingly, behavior from the subordinate assay does not contribute to the treatment effect or syndrome, likely because subordinate focal individuals primarily respond to the dominant fish’s behavior. To our knowledge, this is the first behavioral syndrome to be identified in *A*. *burtoni* at any developmental stage.

Behavior patterns may coalesce into a syndrome due to shared mechanisms (e.g., neuroendocrine regulation), early-life experiences that set individuals along developmentally plastic trajectories, or correlational selection (Bell, 2007; Ketterson and Nolan, Jr., 1999; Stamps, 2003). We found that the behavior of all juveniles was described by the same syndrome, indicating that how the behaviors are related across experimental contexts (i.e., assays) was maintained independently of the early-life social environment. Whether an individual was reared in a group or pair then dictates where along the continuum of the syndrome they fall (Fig 3D).

Pair-reared juveniles appear restricted to one end, whereas group-reared juveniles are represented along the full range of behavioral variation. That there are group-reared juveniles that behaviorally resemble the pair-reared individuals suggests there may be social environments within a group (Saltz et al., 2016) that share key elements with the paired experience. In contrast, the range of possible social roles seems much more restricted in the paired treatment. To identify the causal behavioral and/or sensory cues, it will be necessary to conduct detailed observations of individuals within the rearing environments (Taborsky, 2016). We hypothesize that the complexity of interactions and/or abundance of social sensory cues in groups cause these treatment differences (Taborsky, 2016, e.g., Arnold & Taborsky, 2010). Directly quantifying the range of experience, behavior, and growth within and across early-life environments will be critical to understanding the nature and magnitude of individual phenotypic variation. It can also inform more nuanced selection criteria and analysis methods for comparing focal fish across treatments and tanks than based on size or size ratios alone, as we did in this study.

Activity and social interaction are common components of syndromes in other species, along with bold-shy and proactive-reactive behaviors (Bell, 2007; Conrad et al., 2011; Groothuis and Carere, 2005; Koolhaas et al., 1999; Sih et al., 2004b; Verbeek et al., 1994). For example, large juvenile brown trout are more active and aggressive (Näslund and Johnsson, 2016), similar to our results. Activity-aggression syndromes are also found in a number of other fish species (reviewed in Conrad et al., 2011). For *A*. *burtoni* juveniles, locomotor activity and social interaction may be causally related. First, active individuals may encounter conspecifics more frequently and, as a result, initiate more interactions. Second, juvenile social interactions appear to be prosocial in that they increase the likelihood of future proximity and interaction. In the dominance behavior assay, approaches and displacements for both the focal and subordinate cue fish load in the same direction on PC1. Correlation analysis (data not shown) confirms that, as one member of the pair initiates social interactions, the other member also initiates, potentially leading to more activity. This may be beneficial by increasing shoaling and reducing the risk of predation. Interestingly, adult dominance behavior does not lead to a prosocial response in subordinates, suggesting that although social behavior appears similar across life history stages (Fernald and Hirata, 1979; Fraley and Fernald, 1982), there are important differences.

### Size plays a secondary role in determining juvenile behavioral phenotype

Size is central to understanding the effects of the early-life social environment. Group-reared juveniles were larger than those reared in pairs, which is consistent with previous work showing growth is socially regulated in both juveniles and adults (Fraley and Fernald, 1982; Hofmann et al., 1999). Adult *A*. *burtoni* are also highly sensitive to size during social interactions (Alcazar et al., 2014; Weitekamp & Hofmann, 2017); therefore, size differences could cause differences in behavior. In this study, however, the effect of the early social environment appears larger and more complex than size alone. First, the PCA of behavior from all four assays shows that focal fish SL contributes only moderately to the significant treatment difference for PC1 (Fig 2E), as many other variables load much more strongly on PC1 (i.e., open field, social cue investigation, and dominance behaviors) (see also: Supplemental Fig 5). Second, SL is the strongest contributing variable for PC6, which differs significantly between group- and pair-reared juveniles (Fig 2F). The proportion of the variance described by PC6 (5%) compared to PC1 (43.3%) suggests that size contributes relatively less to the overall treatment effect than behaviors in the open field, social cue investigation, and dominance behavior assays. This is further supported by the finding that in a PCA excluding focal fish SL, PC1 still differs significantly between group- and pair-reared juveniles. In this analysis, PC1 is not associated with SL for either group-or pair-reared juveniles, suggesting size does not drive behavior. The significant, positive association between PC1 and SL for all juveniles results from group-reared juveniles being larger than pair-reared juveniles. Third, SL is also not associated with behavior in any of the four behavior assays (Supplemental Table 2). Finally, the group-reared juveniles that fall within the range of pair-reared juveniles along the continuum of the behavioral syndrome (i.e., high PCA scores, Fig 3) are not the smallest individuals. Together, this evidence suggests that size is secondary in understanding early-life effects on behavior. In future studies, it will be important to test how individual behavior changes over time in relation to both size and developmental stage, which can be decoupled from chronological age in fish (Jonsson and Jonsson, 2014).

### Early-life social experience affects social dynamics when focal juveniles are subordinate

Developmental plasticity can shift behavior in ways that ultimately benefit fitness (Smith and Blumstein, 2008), in part because social behavior has direct consequences for reproductive success (Wilson, 1980, e.g., Henry et al., 2013; Robbins et al., 2007; Young et al., 2006). A majority (64%) of studies show that experimentally increasing the frequency, diversity, or complexity of early-life social experiences enhances social skills or competence (Taborsky, 2016). For example, juvenile *N*. *pulcher* cichlids reared with brood helpers demonstrated more context-appropriate behavior when establishing status, integrating into novel groups, and competing for a resource (Arnold and Taborsky, 2010; Fischer et al., 2015; Taborsky et al., 2013, 2012). We have no evidence yet of an advantage for group-reared juveniles; however, juveniles appear fill the subordinate role differently based on rearing environment, as well as social status experience. While nearly all focal fish successfully established themselves as subordinate (88%) in the assay, and there were no treatment differences in approaches or displacements, there was a significantly larger asymmetry in agonistic efficiency for pair-reared juveniles. There was also a trend for pair-reared juveniles to submit more readily (measured as large fish agonistic efficiency). Status relationships are defined by asymmetrical agonistic displays (Drews, 1993); therefore, pair-reared juveniles may behave more submissively.

We also found that the larger juveniles sampled from the groups and pairs, which we are confident accrued more dominance experience during development given the importance of size for juvenile (and adult, Weitekamp and Hofmann, 2017) social interactions (this study), differed in their patterns of behavior compared to the smaller juveniles. Behaviors from the subordinate assay load on PC2 (16.4% of variance, Fig 2E), and there is a trend for PC2 to differ between the larger and smaller sampled fish (Fig 2B). PC5 (6.6% of variance) differs significantly between the larger and smaller fish. A variety of behaviors load on PC5, including activity in the open field and social cue investigation assays, suggesting that space use is also influenced by status experience and/or relative size within a rearing environment. Overall, the subordinate role is critically important for juveniles because all juveniles will enter adult communities as subordinates. It will be necessary to measure behavior and reproductive success of these juveniles once they are adults in order to determine whether these phenotypes persist or if one is more successful than another (Pradhan, Solomon-Lane, & Grober, 2015).

### Early-life social environment and treatment duration affect neuroendocrine gene expression

We have shown that early-life environments can determine where individuals will fall along the continuum of a newly discovered behavioral syndrome, which raises questions about the underlying mechanisms (e.g., pleiotropic genes and/or neuroendocrine regulation). The behavioral effects we detect as a result of the early-life social environment suggest important variation in the underlying neural regulatory mechanisms. Neuroendocrine stress and sex steroid signaling are likely sites of developmental plasticity in *A*. *burtoni* because they are sensitive to early-life effects (Champagne & Curley, 2005; Shepard et al., 2009), translate environmental conditions and experiences into biological responses (Crespi & Denver, 2005; Wingfield et al., 1990), and regulate behavior (Adkins-Regan, 2009; Solomon-Lane, Crespi, & Grober, 2013). We focused on steroid hormone nuclear receptors, with the addition of CRF, specifically because they regulate the transcription of target genes with a diversity of physiological and behavioral roles (Rochette-Egly, 2005). We found that both the early-life social environment and treatment duration—which corresponds to age, in this study—had a significant effect on gene expression in whole brain. GR1a was the only gene to respond exclusively to treatment, while CRF and GR2 changed significantly over time. Early-life environment and treatment duration interacted to affect the expression of sex steroid hormone receptors ARα and ERα. Finally, although GR1b and MR expression varied across individuals, these genes were not significantly affected by treatment or treatment duration. Factors that we did not measure here (e.g., social status, body size, sex), including individual behavior and position along the behavioral syndrome, are also likely to contribute to important variation in gene expression.

The HPA/I axis has a highly-conserved role in responding to early-life environments (Crespi and Denver, 2005). Our results suggest that developmental plasticity can “tune” the HPI axis in nuanced ways via changes in the density and distribution of different receptors and by affecting circulating glucocorticoid levels (Bernier et al., 2009), over developmental time (e.g., CRF, GR2, Fig 4C, E) and in response to different environments (e.g., GR1a, Fig 4D). Many teleosts, including *A*. *burtoni*, have four glucocorticoid receptors: MR, GR1a, GR1b, and GR2. Receptor 1 has subtypes 1a and 1b, which differ by a nine amino acid insertion between the two zinc fingers in the DNA-binding domain (Bury, 2017; Greenwood et al., 2003; Korzan et al., 2014). These receptors differ substantially in their affinity for cortisol. In adult *A*. *burtoni*, MR is 100-fold more sensitive to cortisol than the GRs and is likely to be occupied with cortisol at basal levels (in fish and tetrapods). GR2 has the next highest sensitivity, followed by GR1a, then GR1b (Arterbery et al., 2011; Bury, 2017; Greenwood et al., 2003). Changes in HPA/I axis function typically manifest as altered baseline levels of circulating glucocorticoids, a higher or lower glucocorticoid ‘peak’ in response to an acute stressor, and/or altered efficiency of the negative feedback loop that returns the system to baseline. Negative feedback, in particular, is regulated by neural GR expression (Bernier et al., 2009; Bury, 2017; Denver, 2009; Kiilerich et al., 2018; Wendelaar Bonga, 1997) and can be affected by early-life experience (Champagne and Curley, 2005; Francis et al., 1999).

Consistent with the distinct roles for different components of the stress axis (Greenwood et al., 2003), our results show differences in expression patterns across HPI axis candidate genes (Fig 4C-G). GR1, specifically, appears to respond to the early-life social environment in *A*. *burtoni* and other teleost species (Fokos et al., 2017; Nyman et al., 2018, 2017; Taborsky et al., 2013). In the group-living cichlid *N*. *pulcher*, for example, increased early-life social complexity led to altered GR1 expression, but not GR2 or MR expression, in whole brain and telencephalon (Nyman et al., 2018, 2017; Taborsky et al., 2013). In *A*. *burtoni*, higher expression of GR1a in group-reared juveniles (Fig 4D) might increase sensitivity to cortisol and result in more efficient negative feedback, making these individuals less susceptible to stress. Alternatively, given that *A*. *burtoni* naturally live in groups (Fernald and Hirata, 1977), paired rearing or isolation may actually decrease efficiency. Juvenile stress physiology should be tested directly because negative feedback mechanisms are complex and involve multiple receptors (Bury, 2017; Kiilerich et al., 2018). Overall, little is known about the differential roles of GR1a and GR1b, and the differences that have been demonstrated appear to be species-specific (Bury, 2017). For *A*. *burtoni*, sensitivity to the early-life social environment may be a defining difference (Fig 4D, F). That GR2 and MR also do not respond to the early environment may be consistent with their roles in baseline glucocorticoid signaling rather than the stress response (Greenwood et al., 2003), although whole brain expression of GR2 (and CRF) is lower adult subordinate males compared to dominants (Chen and Fernald, 2008). Finally, the stress axis undergoes important changes throughout development (Alsop and Vijayan, 2008; Barry et al., 1995; Jeffrey and Gilmour, 2016; Tsalafouta et al., 2018), and lower levels of CRF and GR2 after 5 weeks (Fig 4C, E) could indicate a developmental shift towards lower stress axis activity. Alternatively, familiarity with or predictability of a social environment (e.g., treatment duration) could shift HPI axis function. Future research testing these HPI axis hypotheses promises to uncover important mechanisms of early-life effects on neuroendocrine and behavioral development.

The sex steroid hormone receptors ARα and ERα were unique among our candidate genes in that effect of rearing environment on gene expression was mediated by treatment duration. These genes are also not a part of the HPI axis. For both receptors, expression in pair-reared juveniles was significantly lower after 5 weeks in the rearing environments compared to pair-reared juveniles after 1 week and group-reared juveniles after both 1 week and 5 weeks (Fig 4A, B). Compared to the HPA/I axis, less is known about sex steroid hormones in the context of early-life effects, but there are multiple, non-mutually exclusive mechanisms that could explain these expression patterns. First, it is well-established that the HPI axis interacts with the hypothalamic-pituitary-gonadal axis that regulates reproduction, in part through neural ARs and ERs (Acevedo-Rodriguez et al., 2018; Huffman et al., 2012; Schreck, 2010). While we do not expect HPI axis plasticity to entirely drive the changes in ARα and ERα expression, interactions are likely between these important neuroendocrine axes (see below, Fig 5). Second, early-life social experiences can exert lasting changes in sex steroid hormone receptor expression via epigenetic mechanisms. In rats, for example, ERα in the medial POA is critical to the neuroendocrine regulation of maternal licking and grooming. The rates of maternal care received by female pups subsequently affects their future maternal behavior. The mechanism for this early-life maternal effect has been described in detail and involves brain region-specific epigenetic methylation of the ERα promoter (Cameron et al., 2008). Similar epigenetic mechanisms may regulate ARα and ERα (as well as GR, Turecki and Meaney, 2016) in juvenile *A*. *burtoni*, such that epigenetic marks accrue over time in particular early-life social environments. In our study, expression differences were evident after 5 weeks but not yet after 1 week (Fig 4A, B).

Finally, ARα and ERα are found throughout the social decision-making network, a highly-conserved set of brain regions that, together, are involved in the regulation of social behavior across vertebrates, including *A*. *burtoni* (O’Connell and Hofmann, 2012b, 2011). Sex steroid hormone receptors regulate and respond to social behavior and context (Burmeister et al., 2007; Maruska, 2015; O’Connell and Hofmann, 2012a); therefore, the expression patterns in juveniles could reflect social interactions. ARα and ERα mRNA may increase early in development to facilitate sociality, possibly along with other relevant neuromodulators and receptors. After 5 weeks in pairs, the decrease in transcription (Fig 4A, B) may reflect a less dynamic social decision-making network as a consequence of a social environment that is highly predictable. Conversely, expression for juveniles in groups, the most complex environment in this study, remains high over time. Expression for isolated juveniles, an environment absent of social stimuli, is closest to the expression of pair-reared juveniles after 5 weeks. It is noteworthy that all of the candidate genes show a similar pattern (Fig 4): the expression of pair-reared juveniles after 5 week is the lowest compared to other treatments and time points, and the most similar group is isolated juveniles after 1 week. Future work is needed to understand the functional significance of this downregulation.

An important consideration in interpreting these results, and the co-expression networks below, is that gene expression was measured in whole brain. Although the brain is a heterogeneous tissue made up multiple cell types (e.g., neurons, glia) and regions with distinct functionality, we chose this approach because we did not have an *a priori* expectation as to which brain regions or cell types might be the most critical to examine in our study. We recognize that important variation in gene expression might not be detected using this approach; therefore, future research should use approaches that allow for increased spatial resolution (see below), as well as unbiased (rather than candidate) gene expression analysis (e.g., via RNA-Seq). A genome-scale analysis of expression can provide insight into large numbers of genes simultaneously and suggest novel candidate pathways. However, recent analyses have demonstrated that brain region-specific, or even cell type-specific, gene expression patterns can be inferred from bulk tissue samples (e.g., whole brain) (Kelley et al., 2018).

### Complex co-expression of stress and sex steroid signaling by the early-life social environment

Neuroendocrine systems are dynamic and interact on multiple biological levels (e.g., Acevedo-Rodriguez et al., 2018), including within gene regulatory networks (e.g., Huffman et al., 2012; Korzan, Fernald, & Grone, 2014; O’Connell & Hofmann, 2012); therefore, the expression of other genes can also contribute to the variation in a gene of interest. Based on their co-localization in the POA of *A*. *burtoni* (Korzan et al., 2014), co-localization and correlation in other species (e.g., Meyer & Korz, 2013), and overlapping physiological effects (Crespi & Denver, 2005; Wingfield et al., 1990), the neuroendocrine pathways represented by our candidate genes are likely to functionally interact. We identified striking differences in co-expression networks among juveniles reared in different environments. Expression was highly correlated in pair-reared juveniles (Fig 5A), such that every candidate gene was significantly correlated with at least two others. At the center of the network, ARα shares five significant connections. The two sex steroid hormone genes (ARα, ERα) are also integrated with the stress axis genes, which form distinct smaller networks: CRF-GR1a-GR1b and GR2-MR. In contrast, group-reared juveniles have only one significant partial correlation between ERα and GR1b, a connection that is not present in the pair-reared network (Fig 5B). There are no significant partial correlations for isolated juveniles, suggesting that the neuroendocrine regulatory network is dysregulated, possibly due to isolation acting as a stressor (Galhardo and Oliveira, 2014). These network differences, together with other relevant genes not included in our candidate analysis, might underlie the behavioral differences we identified in the behavioral syndrome, subordinate behavior, or more broadly related to stress response. The differential co-regulation could also serve to make behavior more similar in the face of other neural differences caused by rearing environment, as is the case for some neural sex differences and behavior (De Vries, 2004). These hypotheses can be tested directly using central pharmacological manipulation.

### Integrating the effects of early-life social environments on behavior and brain

Our work demonstrates that early-life social environments shape behavioral phenotype and neuroendocrine gene expression in powerful ways for *A*. *burtoni* juveniles. In the present study, we quantified behavior and gene expression in separate experiments in order to focus on different developmental time points. Taken together, our results allow us to generate strong hypotheses about the mechanisms, consequences, and developmental time course of early-life social effects. By focusing on early time points for gene expression—after 1 and 5 weeks in the social environments—we aimed to identify highly sensitive components of these neuroendocrine systems. Based on our results, we expect the HPI axis via GR1a expression to respond rapidly to different social environments. In contrast, the treatment differences in ARα and ERα emerge over time. After 8-10 weeks, when we analyzed behavior, we hypothesize that altered stress physiology aligns with the behavior patterns sensitive to early-life effects, together forming a specific kind of syndrome called a coping style. Individuals range in coping style from proactive to reactive, such that proactive copers are more active, aggressive, and less responsive to stress (i.e., lower baseline glucocorticoid levels, faster negative feedback) than reactive copers (Koolhaas et al., 1999). As juveniles approach reproductive maturity (as early as 12 weeks old, Fraley and Fernald, 1982), we expect social environment and age to continue to interact for ARα and ERα. Maturation is socially regulated, including by the early-life social environment; therefore, we predict that expression will reflect ongoing social demands, as well as emerging reproductive behavior (Fraley and Fernald, 1982). This work can begin to address the fact that across species, remarkably little is known about the mechanisms that shape the ontogeny of behavior (Taborsky, 2016).

Understanding the full scope and consequences of early-life effects ultimately requires measuring brain and behavior in the same individuals, throughout development and into adulthood. Our results suggest that brain regions that express GR1a, ARα, and ERα (Greenwood et al., 2003; Korzan et al., 2014; Munchrath and Hofmann, 2010), along with brain regions of the social decision-making network (O’Connell and Hofmann, 2012b), are likely to be sensitive to early-life effects and could cause the observed changes in behavior. Interestingly, the POA—a critical node in the social decision-making network (O’Connell and Hofmann, 2012b)—contains GR1a, GR1b, GR2, MR, ARα, and ERα in adult *A*. *burtoni* (Korzan et al., 2014; Munchrath and Hofmann, 2010). Additional nodes, such as the hippocampus and amygdala are also likely sites of overlap. These brain regions are important in spatial cognition and emotional processing, respectively, and are central to HPI axis negative feedback (Denver, 2009). Interactions between the HPI axis and sex steroid hormone signaling, including in the POA, could be a mechanism for the social regulation of development (Fraley and Fernald, 1982; Korzan et al., 2014; Solomon-Lane et al., 2013; Wada, 2008). Overall, this research can uncover the neuroendocrine mechanisms by which early-life social experience gives rise to individual variation in adults, which is critical to understanding subsequent disparities in fitness and health.

## Supporting information

## Acknowledgements

We thank Savannah Clapp, Pamela Del Valle, and Najah Hussain for assistance with data collection and fish maintenance and care. We thank Dr. Becca Young for helpful comments on earlier versions of this manuscript and members of the Hofmann Lab for discussion and feedback.

