## Supplementary material for "Early-life social environment alters juvenile behavior and neuroendocrine function in a highly social cichlid fish"

**Supplemental Table 1:** Primers used for quantitative real-time PCR analyses.

| Gene | Primer sequence (forward) | Primer sequence (reverse) |
|---|---|---|
| ARα | CAG GAA TGC CGC TGT ATC TG | TGA GGA ATC GCA CTT GGG TA |
| ERα | CTA CGA AGT GGG CAT GAT GAA A | GGT CTT TGG CTG GTT TGT CTC T |
| GR2 | TGC CTC TGT CAC TGC CAC CGT AG | AGT CGT CTG CGT CTG AAG TAA CTG |
| GR1a | TCA TAA GAT CTG TTT GGT GTG CTC | GTA GTT GTG CTG GCC TTC AAC |
| GR1b | TGT TGG CTT CTC CGG TTC ATC AC | GTT GTG CTG GCC ATC TGT GTT T |
| MR | CGT TAA TGG AGT CGT GGA AAT C | GAG GAC GGT TGT CTC AGT GG |
| CRF | CGA ACT CTT TCC CAT CAA CGT CCA | AGC GCC CTG ATG TTC CCA ACT TTA |
| 18S | CCC TTC AAA CCC TCT TAC CC | CCA CCG CTA AGA GTC GTA TT |
| G3PDH | CAC ACA AGC CCA ACC CAT AGT CAT | CCA CCG CTA AGA GTC GTA TT |

**Supplemental Table 2:** Linear regressions of focal fish standard length with behavior in the open field, social cue investigation, dominance, and subordinate behavior assays for group-reared and pair-reared juveniles separately. The zones of the tank refer to the frequency of entering that zone. Adjusted $R^2$ values are reported. None of the results is significant following a false discovery rate correction.

| | | Group-reared | | Pair-reared | |
|---|---|---|---|---|---|
| **Assay** | **Behavior** | **r-squared** | **p-value** | **r-squared** | **p-value** |
| **Open field** | Investigate zone | 0.02 | 0.23 | 0.16 | 0.058 |
| | Far zone | 0.03 | 0.21 | 0.13 | 0.084 |
| | Close zone | 0.0049 | 0.30 | 0.12 | 0.087 |
| | Territory zone | 0.047 | 0.16 | 0.20 | 0.036 |
| **Social cue investigation** | Investigate zone | -0.045 | 0.94 | -0.037 | 0.54 |
| | Far zone | -0.044 | 0.8573 | -0.0096 | 0.37 |
| | Close zone | 0.051 | 0.15 | 0.12 | 0.086 |
| | Territory zone | 0.12 | 0.052 | 0.11 | 0.099 |
| **Dominance behavior** | Approach | 0.014 | 0.26 | -0.032 | 0.50 |
| | Displace | 0.037 | 0.18 | -0.052 | 0.70 |
| | Territory zone (focal) | -0.020 | 0.47 | 0.23 | 0.027 |
| | Territory zone (subordinate fish) | 0.018 | 0.25 | 0.15 | 0.065 |
| | Territory zone (both) | 0.015 | 0.26 | 0.10 | 0.11 |
| **Subordinate behavior** | Approach | -0.038 | 0.70 | -0.038 | 0.55 |
| | Displace | -0.035 | 0.63 | -0.057 | 0.78 |
| | Submit | -0.0019 | 0.34 | -0.051 | 0.68 |
| | Territory zone (focal) | 0.074 | 0.11 | -0.044 | 0.60 |
| | Territory zone (subordinate fish) | 0.21 | 0.013 | 0.041 | 0.21 |
| | Territory zone (both) | 0.039 | 0.18 | 0.091 | 0.12 |

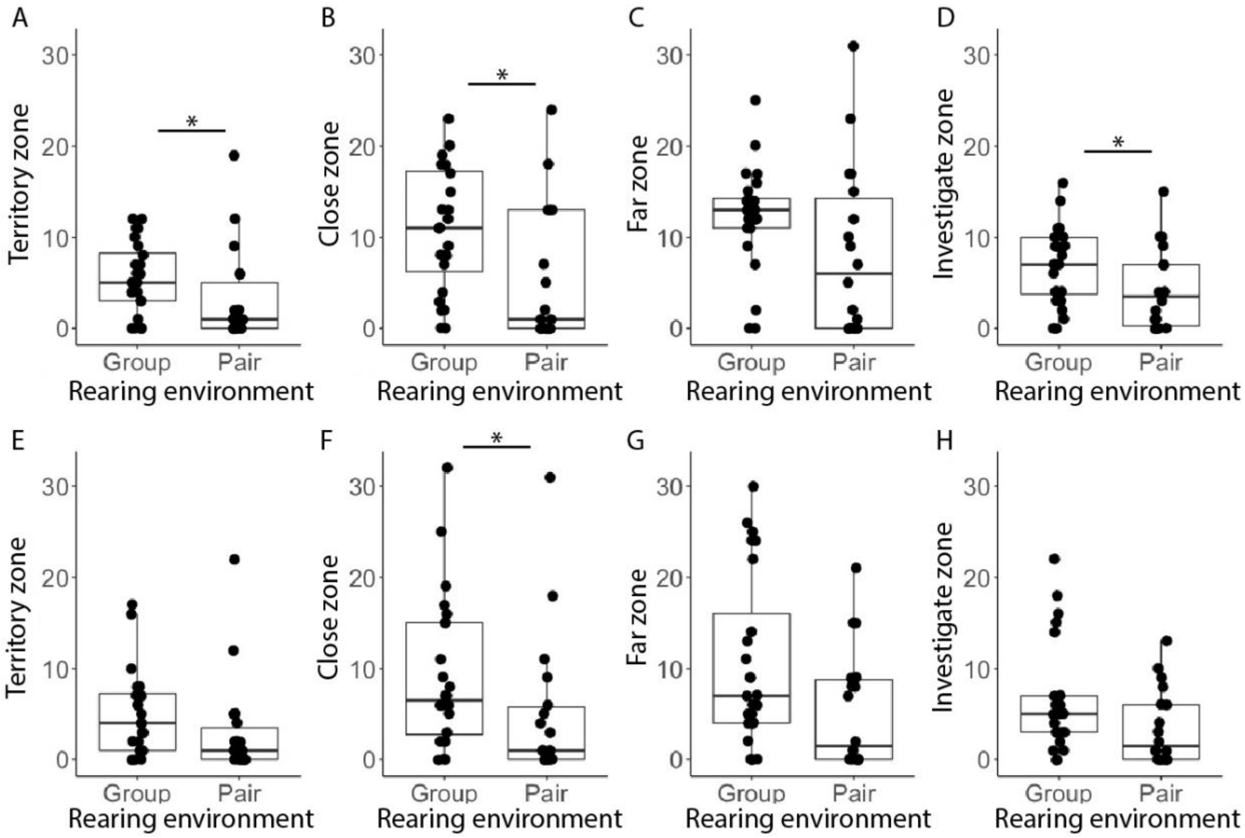


**Supplemental Figure 1:** Frequency of focal fish reared in groups and pairs entering each territory zone during the open field assay (A-D) and social cue investigation assay (E-H). Each assay lasted for 10 min. See Figure 1 for a map of the tank zones. *p<0.05.

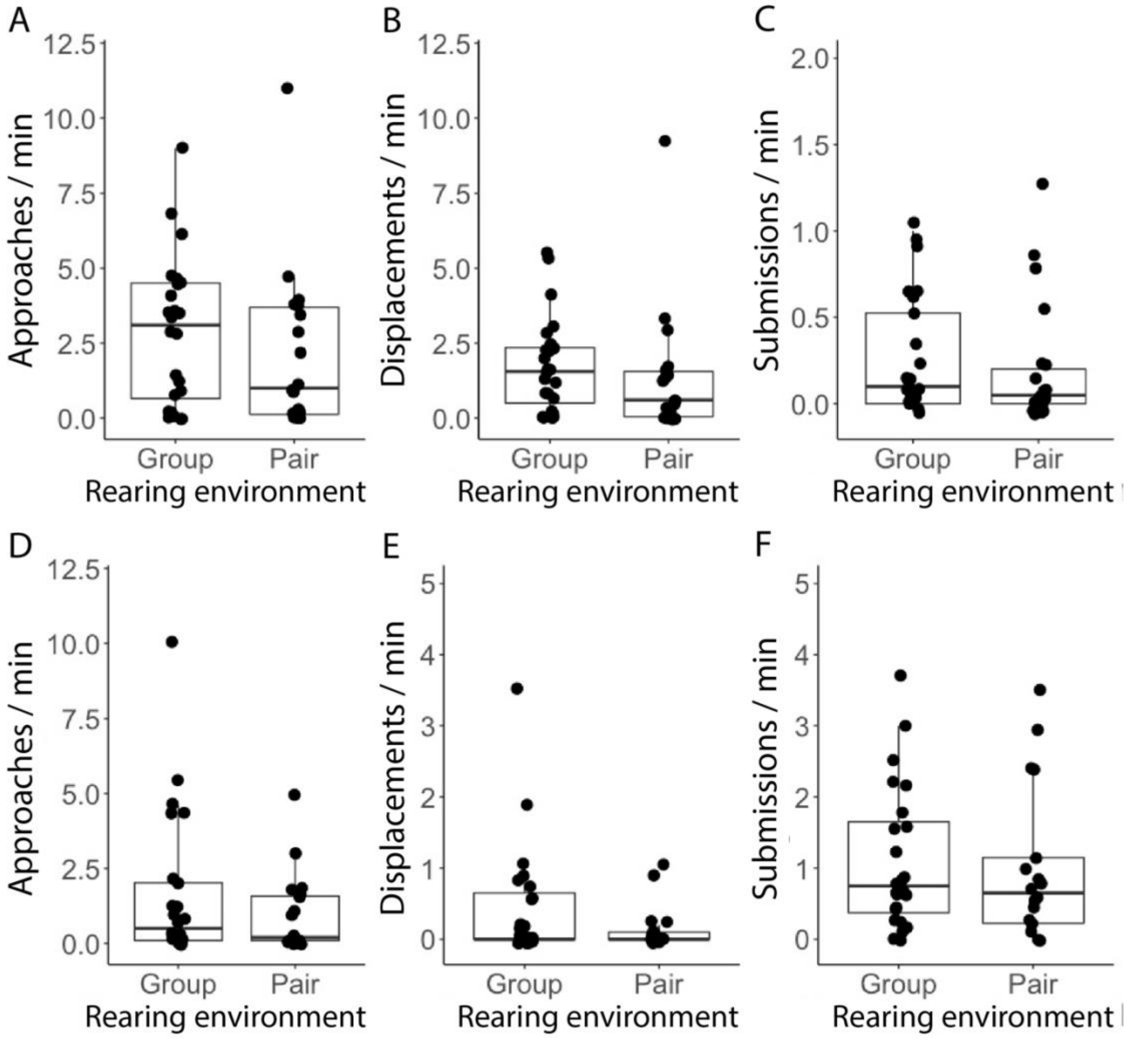


**Supplemental Figure 2:** Group- and pair-reared focal fish social behavior in the dominance (A-C) and subordinate (D-F) assay.

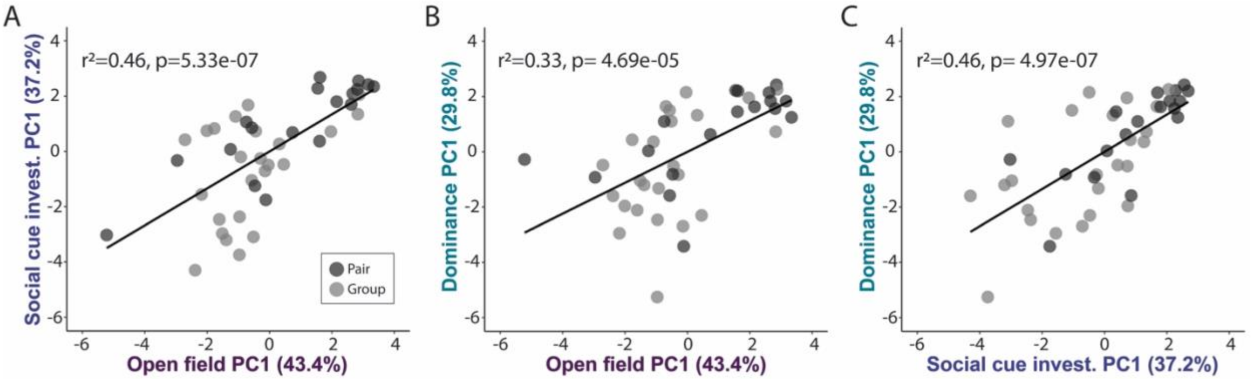


**Supplemental Figure 3:** Separate principal component analyses of open field, social cue investigation, and dominance behavior, including focal and non-focal fish variables (behavior, size). For each assay, PC1 differs significantly between group- and pair-reared juveniles. Linear regression analysis shows that PC1s across tests are positively associated: open field x social cue investigation (A), open field x dominance behavior (B), and social cue investigation x dominance behavior (C). Percentages refer to the amount of variation explained by that component. Pair (n=18 individuals). Group (n=24 individuals).

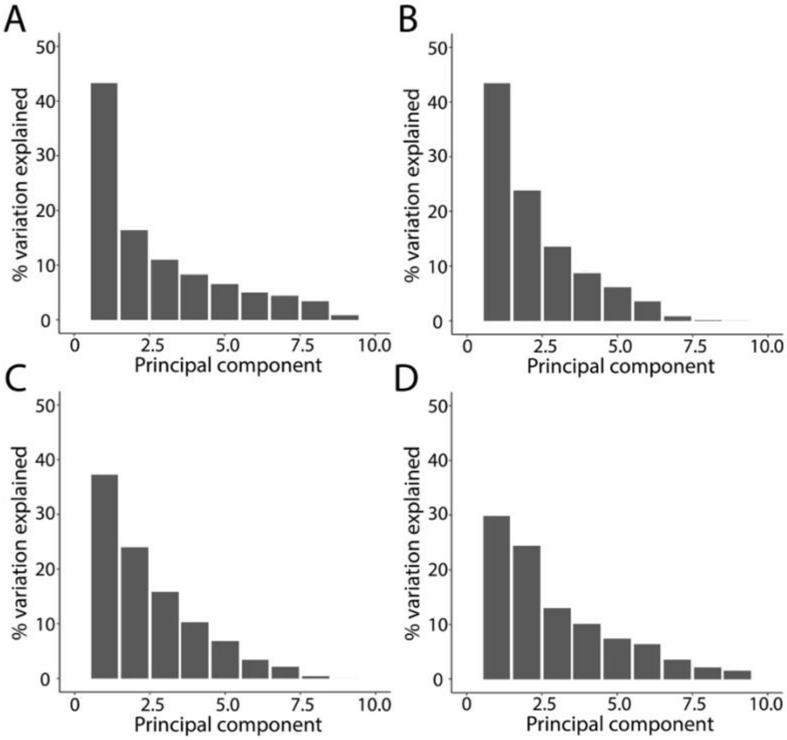


**Supplemental Figure 4:** The percentage of the variation explained by principal components for analyses including all four behavior assays (open field, social cue investigation, dominance, and subordinate behavior, A), the open field test (B), the social cue investigation (C), and dominance behavior (D).

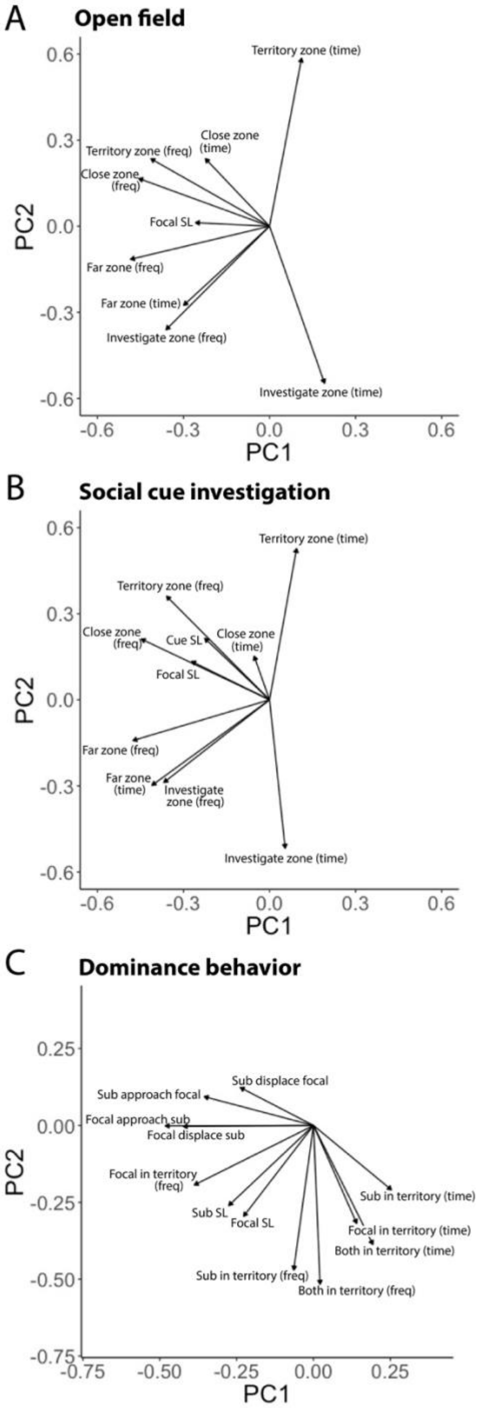


**Supplemental Figure 5:** Vector plots of principal component 1 x principal component 2 for principal components analysis of open field (A), social cue investigation (B), and dominance behavior (C) assays. For each assay, PC1 differ significantly between group- and pair-reared juveniles. There is no effect for PC2.

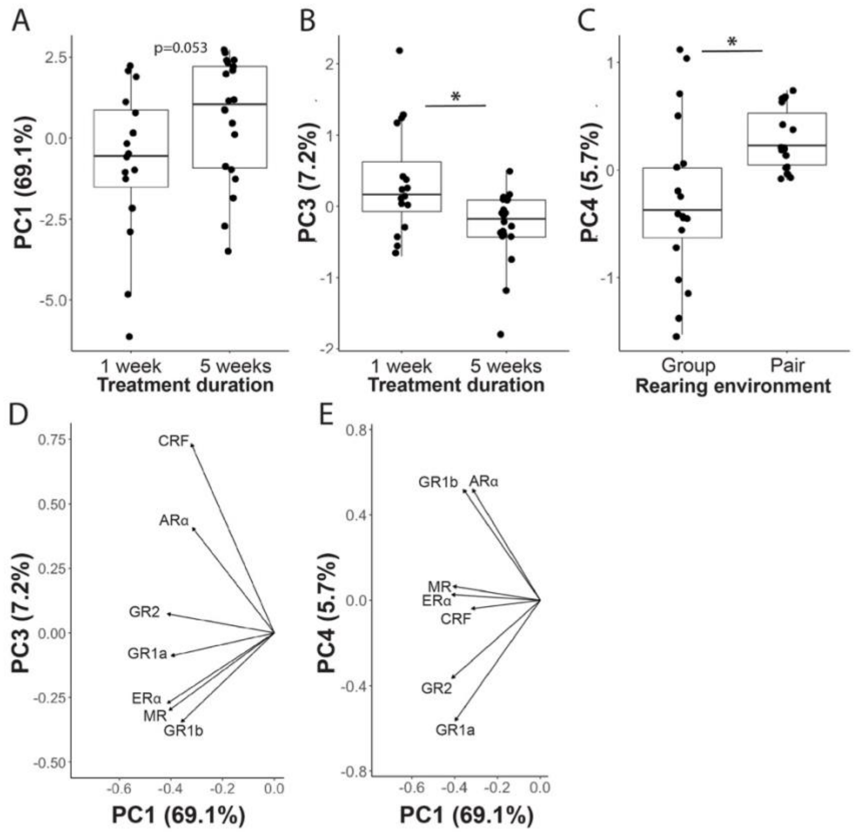


**Supplemental Figure 6:** To gain a more holistic understanding of how rearing environment and/or treatment duration affect variation in neuroendocrine gene expression, we used principal component analysis. T-tests, or Mann-Whitney-Wilcoxon tests, if appropriate, were used to compare group- and pair-reared juveniles and expression following 1 vs. 5 weeks in rearing environments. Dunn's test was used for *post hoc* analysis of significant results. PC1 accounts for 69.1% of the variance in the data. While there were no differences in PC1 based on rearing environment (W=113, p=0.13), there was a trend for differences based on treatment duration (W=99, p=0.053; A). PC3 (7.17% of the variance) differed significantly between juveniles in treatment groups for 1 vs. 5 weeks (W=234, p=0.018, d=0.91; B). PC4 (5.7% of the variance) differed significantly between group- and pair-reared juveniles (t=2.80, p=0.011, d=0.93; C). Vector plots show gene loadings on PC1 and PC3 (D) and PC1 and PC4 (E).
